# μ opioid receptors acutely regulate adenosine signaling in a thalamo-striatal circuit

**DOI:** 10.1101/2021.05.25.445648

**Authors:** Sweta Adhikary, William T. Birdsong

## Abstract

Endogenous adenosine plays a crucial role in maintaining energy homeostasis and adenosine levels are tightly regulated across neural circuits. In the dorsal medial striatum (DMS) adenosine inhibits neurotransmitter release, but the source and mechanism underlying its accumulation are largely unknown. Opioids also inhibit neurotransmitter release in the DMS and influences adenosine accumulation after prolonged exposure. However, how these two neurotransmitter systems interact acutely is also largely unknown. This study demonstrates that activation of μ opioid receptors (MORs), but not δ opioid receptors (DORs) or κ opioid receptors (KORs), inhibits tonic activation of adenosine A_1_Rs via a cyclic adenosine monophosphate (cAMP) dependent mechanism in both male and female mice. Further, selectively knocking-out MORs from presynaptic terminals and postsynaptic medium spiny neurons (MSNs) revealed that activation of MORs on D_1_R positive MSNs, but not D_2_R positive MSNs, is necessary to inhibit tonic adenosine signaling on presynaptic terminals. Given the role of D_1_R positive MSNs in movement and motivated behaviors, these findings reveal a novel mechanism by which these neurons regulate their own synaptic inputs.

**Significance Statement:** Understanding interactions between neuromodulatory systems within brain circuits is a fundamental question in neuroscience. The present work uncovers a novel role of opioids in acutely inhibiting adenosine accumulation and subsequent adenosine receptor signaling in the striatum by inhibiting the production of cAMP. Adenosine receptor signaling regulates striatal neurotransmitters including glutamate, GABA, dopamine and acetylcholine. Furthermore, interactions between adenosine_2A_ receptors and numerous other GPCRs, including D_2_ dopamine and CB_1_ cannabinoid receptors, suggest that endogenous adenosine broadly modulates striatal GPCR signaling. Additionally, this work discovered that resting endogenous adenosine is released by D_1_, but not D_2_ receptor positive MSNs, suggesting that opioid signaling and manipulation of D_1_R-expressing MSN cAMP activity can broadly affect striatal function and behavior.

## Introduction

Opioids such as morphine acutely mediate analgesia and long-term use leads to dependence and potentially addiction. The thalamus and dorsal medial striatum are important for regulating opioid dependence and modulating goal-directed behavior respectively (Balleine, Delgado, and Hikosaka 2007; Zhu et al. 2016). Opioids are known to inhibit both excitatory input to the striatum, and local GABA release within the striatal micro-circuitry (Atwood, Kupferschmidt, and Lovinger 2014; Banghart et al. 2015; Birdsong et al. 2019). Additionally, agonists selective to the adenosine A_1_Rs also inhibit glutamate release in the striatum (Brundege and Williams 2002) and influence striatal dynamics. Thus, understanding the role of opioid receptors and A_1_Rs in modulating excitatory inputs to the striatum, and the potential interaction between these receptors, is important to understand how multiple neurotransmitter systems influence striatal activity.

Morphine binding to MOR activates G_i/o_ heterotrimeric G-proteins to inhibit adenyl cyclase (AC) and consequently decreases cAMP levels (Heijna et al. 1992; Izenwasser, Buzas, and Cox 1993). Acutely, this inhibition of cAMP, along with other effectors, ultimately inhibits neuronal activity and neurotransmitter release. Similarly, activation of the A_1_Rs also inhibits AC, and under basal conditions there is a resting extracellular adenosine tone in the striatum. This resting adenosine tone can tonically activate A_1_Rs, inhibiting neurotransmitter release (Brundege and Williams 2002). The fact that adenosine and opioids both act through the same effector systems suggests that these two neurotransmitters can influence each other’s signaling. But neither the role of opioids in modulating resting adenosine levels nor the source of this resting adenosine tone is known.

Agonists selective to MOR, but not DOR, potently inhibit glutamate release from thalamus onto the striatum (Birdsong et al. 2019; Muñoz, Haggerty, and Atwood 2020), but the role of KORs in this circuit has not been examined. However, there is evidence that MOR, DOR and KOR are widely expressed throughout the striatum (Al-Hasani et al. 2015; Banghart et al. 2015; Massaly et al. 2019; Mansour et al. 1994; Muschamp and Carlezon 2013; Nestler and Carlezon 2006) and have been shown to inhibit neurotransmitter release in the striatum to varying degrees in a synapse-specific manner (Tejeda et al. 2017). Therefore, although all three subtypes of opioid receptors are present in the striatum, they potentially modulate the activity of the striatum and interact with A_1_R signaling uniquely.

The present study examines the functional interaction between opioid receptors and adenosine signaling, the mechanism underlying extracellular adenosine accumulation, and the source of adenosine release in the striatum using a combination of brain slice electrophysiology, pharmacology, optogenetics, and genetic manipulation of MOR expression in mice. Optically-induced excitatory post synaptic current (oEPSCs) were recorded in striatal medium spiny neurons following optical excitation of channelrhodopsin-expressing medial thalamic axon terminals in the dorsomedial striatum. The facilitation of oEPSC amplitude by the A_1_R antagonist 8-cyclopentyl-1,3-dipropylxanthine (DPCPX) was used to measure tonic A_1_R activation and as a proxy for extracellular adenosine accumulation. The results show (1) morphine inhibits tonic adenosine accumulation by inhibiting cAMP, (2) this inhibition is specific to MOR agonists and not DOR or KOR agonists, and (3) MOR regulation of dorsomedial striatal adenosine levels requires MOR expression on D_1_R positive MSNs.

## Materials and Method

### Animals

Male and female C57BL/6J mice (8–10 weeks old) were bred in house and were housed under a 12-hr-light/dark cycle. Food and water were available *ad libitum*. Mice with exons 2 and 3 of the oprm1 gene flanked by the LoxP cassette (FloxedMor; *Oprm1*^*fl/fl*^; JAX stock #030074), with a genetic background of 75:25% of C57BL/6J were provided by Dr. Brigitte L. Kieffer. Vglut_2_-*cre* mice (*Slc17a6*^*tm2(cre)Lowl*^; JAX stock #016963) were purchased from the Jackson Laboratory. The two mice were crossed to generate FloxMor-Vglut_2_-*cre* mice that lack MORs in presynaptic terminals. A_2A_-*cre* mice (*Tg(Adora2a-cre)KG139Gsat*; MMRC stock #036158-UCD) were provided by Dr. Tianyi Mao and were crossed with FloxedMor mice to generate FloxedMor-A_2A_-*cre* mice lacking MORs in D_2_ positive MSNs. D_1_-*cre* mice (*Tg(Drd1-cre)EY262Gsat*; MMRC stock #030989-UCD) were provided by Dr. Christopher Ford and were crossed with FloxedMOR mice to generate FloxedMOR-D_1_-*cre* mice lacking MORs in D_1_ positive MSNs. All animal experiments were conducted in accordance with the National Institutes of Health guidelines and with approval from the Institutional Animal Care and Use Committee of the Oregon Health & Science University (Portland, OR).

### Viral injection

Mice were anesthetized with isoflurane and placed in a stereotaxic frame (Kopf Instruments, Tujunga, CA, with custom modifications) for microinjections of recombinant adeno-associated virus (AAV2-syn-CsChR-GFP) to express channelrhodopsin. A glass pipette filled with 40 nL of virus was injected into the medial thalamus (Nanoject II, Drummond Scientific, Broomall, PA; BCJ: custom-built injector based on a MO-10, Narishige, Amityville, NY). Injection coordinates for MD are in mm for medial/lateral (M/L), anterior/posterior from bregma (A/P), and dorsal/ventral from the top of the skull directly over the target area: M/L: +/-0.55, A/P: -1.2, D/V: -3.4. Electrophysiology experiments were done 2-3 weeks after viral injections.

### Drugs

Morphine sulfate was obtained from the National Institute on Drug Abuse (Baltimore, MD). Naloxone and dizocilpine maleate (MK801) were from Abcam (Cambridge, MA). 8-Cyclopentyl-1,3-dipropylxanthine DPCPX, CGS21680, SKF81297, Mecamylamine, CGP 55845, and MPEP were from Tocris Bioscience (Ellisville, MO). Scopolamine, Adenosine, [Met^5^] Enkephalin (ME), Bestatin, Thiorphan, and R0-20-1724 were from Sigma Aldrich (St. Louis, MO). Picrotoxin was from Hello Bio. ME, Morphine, Adenosine, Naloxone, MPEP, Scopolamine, and Mecamylamine were dissolved in water, diluted in artificial cerebrospinal fluid (ACSF) and applied by bath superperfusion. Bath perfusion of ME was with bestatin (10 μM) and thiorphan (1 μM) to limit breakdown of ME. Picrotoxin was directly dissolved in ACSF. DPCPX, CGS21980, SKF81297 and R0-230853 were dissolved in dimethyl sulfoxide (DMSO), diluted in ACSF and applied during incubation and by bath superperfusion.

### Tissue Preparation

Acute brain slice preparation was performed as previously described (Birdsong et al. 2019). Briefly, mice were deeply anesthetized and euthanized using isoflurane. Brains were removed, blocked, and mounted in a vibratome chamber (VT 1200S; Leica, Nussloch, Germany). Coronal slices (242 μM) were prepared in warm (34°C) ACSF containing (in millimolars) 126 NaCl, 2.5 KCl, 1.2 MgCl2, 2.6 CaCl2, 1.2 NaH2PO4, 21.4 NAHCO3, and 11 D-glucose with +MK-801 (10 mM). Slices were allowed to recover in warm ACSF containing +MK-801 (10 µM) for at least 30 minutes and then stored in glass vials at room temperature with oxygenated (95% O2/ 5% CO2) ACSF until used.

### Brain slice electrophysiology

Slices were hemisected and then transferred to the recording chamber, which was continuously superfused with 34°C carbogenated ACSF at 1.5–2 ml/min with (in µM): 0.2 GABA_B_-receptor antagonist CGP 55845, 10 GABA_A_-receptor antagonist picrotoxin, one nicotinic acetylcholine receptor antagonist mecamylamine, 0.1 muscarinic acetylcholine receptor antagonist scopolamine and 0.3 metabotropic glutamate receptor five antagonist MPEP. Whole-cell recordings from medium spiny neurons (MSNs) in the dorsal medial striatum were obtained with an Axopatch 200A amplifier (Axon Instruments) in voltage-clamp mode, holding potential (V_hold_ = -75 mV). Recording pipettes (Sutter Instruments, Novato, CA) with a resistance of 2.8–3.5 MΩ were filled with an internal solution of (in millimolars) 110 potassium gluconate, 10 KCl, 15 NaCl, 1.5 MgCl_2_, 10 HEPES, 1 EGTA, 2 Na_2_ATP, 0.3 Na_2_GTP, 7.8 phosphocreatine; pH 7.35– 7.40 ∼280 mOsm). Data were filtered at 10 kHz and collected at 20 kHz with AxographX. Only recordings in which the series resistance remained < 18 MΩ or changed by less than 20 percent throughout the experiment were included. A TTL-controlled LED driver and 470 nm LED (Thorlabs, Newton, NJ) was used to illuminate the slice through the microscope objective directly over the recorded cell with ∼1 mW of power for 0.5 ms or 1 ms.

### Electrophysiology data analysis

Data were analyzed in Axograph. Peak current amplitude was calculated relative to mean current during 50 ms baseline prior to the stimulus. Statistical analysis was performed using GraphPad Prism 8 software (GraphPad Software Inc., San Diego, CA). For the time course and summary data, baseline oEPSCs were normalized to oEPSCs amplitudes three to four minutes prior to baseline (prebaseline condition, not shown). All other conditions were normalized to oEPSC amplitudes three to four minutes before drug application. Summary data were presented as the averages of six to 10 trials beginning three to four minutes after drug application after steady state was achieved. For all conditions, mice were used to obtain at least five technical replicates per group; if more than six could be analyzed, all were included. Values are presented as average +/-SEM. Statistical analysis was performed on normalized data. Statistical comparisons were made with paired ratio T-test, one-way repeated measures ANOVA, or one-way ANOVA, followed by multiple comparison adjusted Tukey’s post hoc tests. For all experiments, P <0.05 was used to describe statistical significance.

## Results

### Thalamo-striatal glutamate release is sensitive to both opioid and adenosine agonists

Adeno-associated virus (AAV) type 2 encoding channelrhodopsin was microinjected into the medial thalamus, and whole-cell voltage-clamp recordings were made from medium spiny neurons in the dorsal medial striatum (DMS) (Fig 1A). Striatal MSNs were identified by their hyperpolarized resting membrane potential, low input resistance and a long delay to the initial spike (Kreitzer 2009). Glutamate release was evoked by optical stimulation with 470-nm light, and AMPA receptor-mediated excitatory postsynaptic currents (oEPSCs) were pharmacologically isolated and recorded as previously described (Birdsong et al. 2019). After a stable baseline of oEPSCs was established, partial agonist morphine (1 μM) was superperfused, followed by antagonist naloxone (1 μM) (Fig 1C, E). The inhibition by morphine was determined by averaging the oEPSC three to five minutes after drug perfusion and normalizing the response to the average of oEPCS three to five minutes before drug perfusion. Morphine decreased the amplitude of the oEPSCs and this inhibition was reversed by naloxone (Fig 1C, E, F; morphine: 0.80 ± 0.05 fraction of baseline, p = 0.0002; naloxone: 0.98 ± 0.01 fraction of baseline, p = 0.002, n = 8 cells, 4 mice, F_(2, 14)_ = 17.29, one-way repeated measures ANOVA, Tukey’s multiple comparisons test). In separate cells, an A_1_R agonist cyclopentyladenosine (CPA, 1 μM) was superperfused, followed by antagonist DPCPX, 200 nM), (Fig D, E). CPA decreased the amplitude of the oEPSCs and this inhibition was reversed by DPCPX (Fig 1D, E, G; CPA 0.37 ± 0.04 fraction of baseline, p < 0.0001; DPCPX: 1.3 ± 0.08 fraction of baseline, p < 0.0001 n = 7 cells, 6 mice, F_(2, 12)_ = 87.95, one-way repeated measures ANOVA, Tukey’s multiple comparisons test). Additionally, DPCPX caused a significant over-reversal of the amplitude of the oEPSCs (Fig 1D, E, G), suggesting tonic inhibition of glutamate release by activation of A_1_Rs that was blocked by DPCPX. Thus, glutamate release in the thalamo-striatal synapses is regulated by both MORs and A_1_ receptors, and there is an additional tonic activation of A_1_Rs by endogenous adenosine.

**Figure 1.**
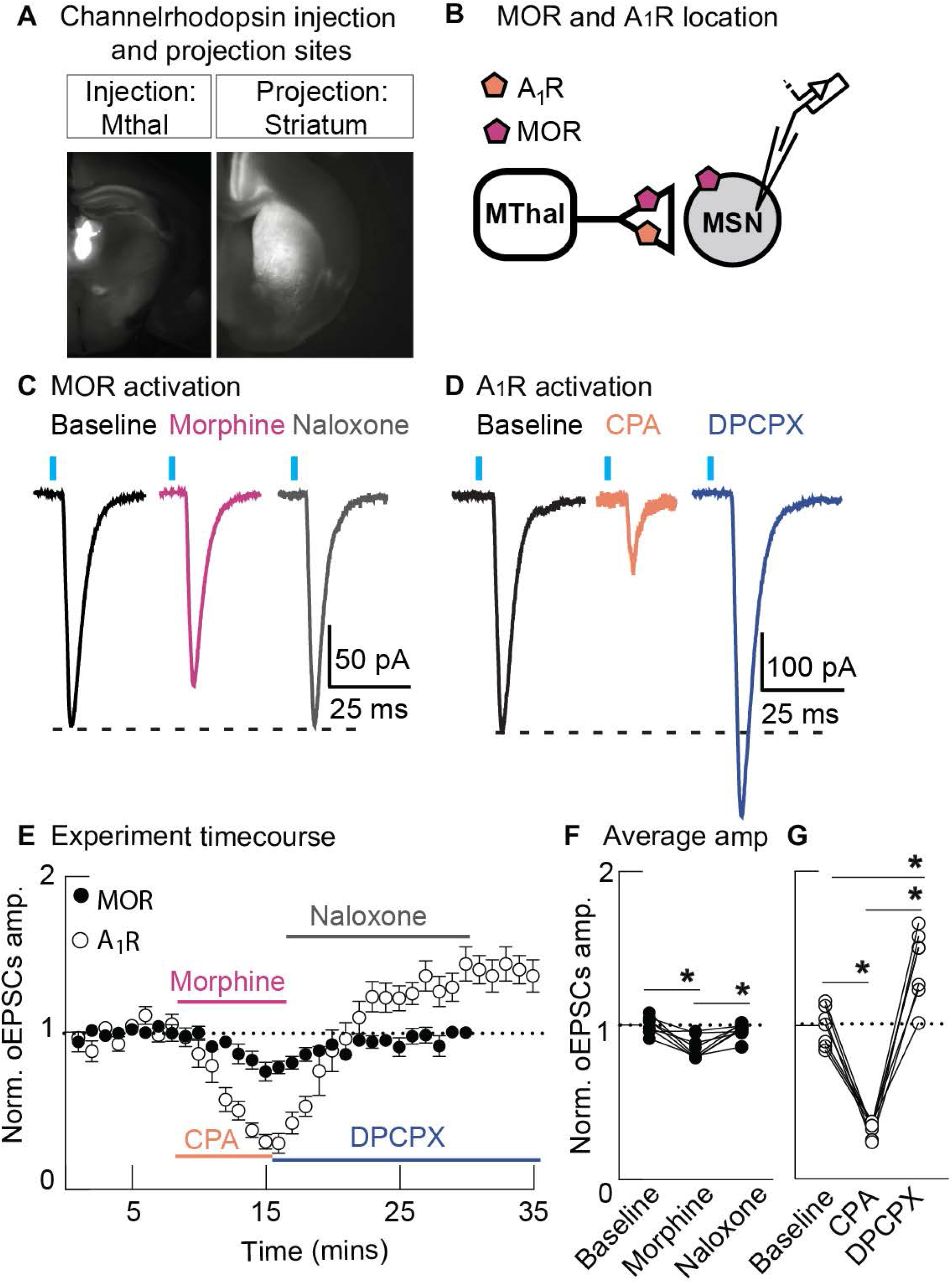
Activation of both μ opioid receptor and adenosine A_1_ receptor leads to inhibition of thalamo-striatal oEPSCs. **(A)** An acute mouse brain slice example of overlaid brightfield and epifluorescent images showing the viral injection site (Mthal; left) and the axonal projections (Striatum; right). **(B)** Schematic showing the locations of both A_1_Rs and MORs in the thalamo-striatal synapse. **(C)** Representative oEPSCs evoked by 470 nm light (black label), inhibition of oEPSC amplitude by morphine (1 μM; pink label), and reversal by naloxone (1 μM; gray label). **(D)** Representative oEPSCs evoked by 470 nm light (black label), inhibition of oEPSC amplitude by CPA (1 μM; orange label), and over reversal by DPCPX (200 nM; blue label). **(E)** Plot of the time course of normalized oEPSC amplitude for cells treated with morphine, followed by naloxone (dark circles; n = 8 cells, 4 mice), and for cells treated with CPA, followed by DPCPX (clear circles; n = 7 cells, 6 mice). **(F)** Mean summary data of normalized oEPSC amplitude in baseline condition, after morphine perfusion, followed by naloxone (morphine: 0.80 ± 0.05 fraction of baseline, p = 0.0002; naloxone: 0.98 ± 0.01 fraction of baseline, p = 0.002, n = 8 cells, 4 mice, F_(2, 14)_ = 17.29, one-way repeated measures ANOVA, Tukey’s multiple comparisons test). **(G)** Mean summary data of normalized oEPSC amplitude in baseline condition, after CPA perfusion, followed by DPCPX (CPA 0.37 ± 0.04 fraction of baseline, p < 0.0001; DPCPX: 1.3 ± 0.08 fraction of baseline, p < 0.0001 n = 7 cells, 6 mice, F_(2, 12)_ = 87.95, one-way repeated measures ANOVA, Tukey’s multiple comparisons test). Line and error bars represent mean ± SEM; ^*^ denotes statistical significance.

### μ opioid receptor regulation of tonic adenosine A_1_ receptor activation

Since both MORs and A_1_Rs are coupled to G_i/o_ G-proteins and both are present in thalamic terminals, functional interaction between the two receptors in regulating glutamate release was examined. oEPSCs were evoked as described above and DPCPX (200 nM) was superperfused to measure the effect of tonic A_1_R activation. DPCPX increased oEPSC amplitude (Fig 2A, C, D; DPCPX: 1.3 ± 0.06 fraction of baseline, p = 0.0003, n = 10 cells, 7 mice, t(9) = 5.752, ratio paired T-test). In separate cells, morphine (1 μM) was superperfused, followed by DPCPX. Morphine reduced the amplitude of oEPSCs (Fig 2B, C, E,) as expected. However, in the presence of morphine, DPCPX did not increase oEPSC amplitude (Fig 2B, C, E; morphine 0.78 ± 0.03 fraction of baseline, p = 0.0011; DPCPX: 0.77 ± 0.05 fraction of baseline p = 0.04, and 0.99 ± 0.06 fraction of morphine, p = 0.84, n = 6 cells, 4 mice, F_(2, 10)_ = 14.00, one-way repeated measures ANOVA, Tukey’s multiple comparisons test), suggesting that morphine inhibited the tonic activation of A_1_Rs. To determine whether morphine inhibited the tonic A_1_R activation through MOR activation, or a non-specific morphine effect, global MOR knockout (KO) mice were used. Similar to WT mice, DPCPX increased the oEPSC amplitude in slices from these mice (Fig 2F, H, I; DPCPX: 1.37 ± 0.05 fraction of baseline, p = 0.0001, n = 8 cells, 5 mice, t(7) = 8.273, ratio paired T-test). In contrast to WT mice, morphine did not reduce the amplitude of oEPSCs (Fig 2G, H, J: morphine: 1.0 ± 0.05 fraction of baseline, p = 0.9863, n = 6 cells, 3 mice) in slices from MOR KO mice. Further, in the presence of morphine, DPCPX increased oEPSC amplitude (Fig 2G, H, J; DPCPX: 1.4 ± 0.07 fraction of baseline, p = 0.0006, 1.3 ± 0.09 fraction morphine, p = 0.0008, n = 6 cells, 3 mice, F_(2, 12)_ = 17.46, one-way repeated measures ANOVA, Tukey’s multiple comparisons test). There was no difference in the increase in oEPSC amplitude between control slices and slices in morphine, suggesting that MORs are required for morphine to modulate tonic adenosine levels. Therefore, morphine inhibits the tonic activation of A_1_Rs in the thalamo-striatal circuit by activating MORs.

**Figure 2.**
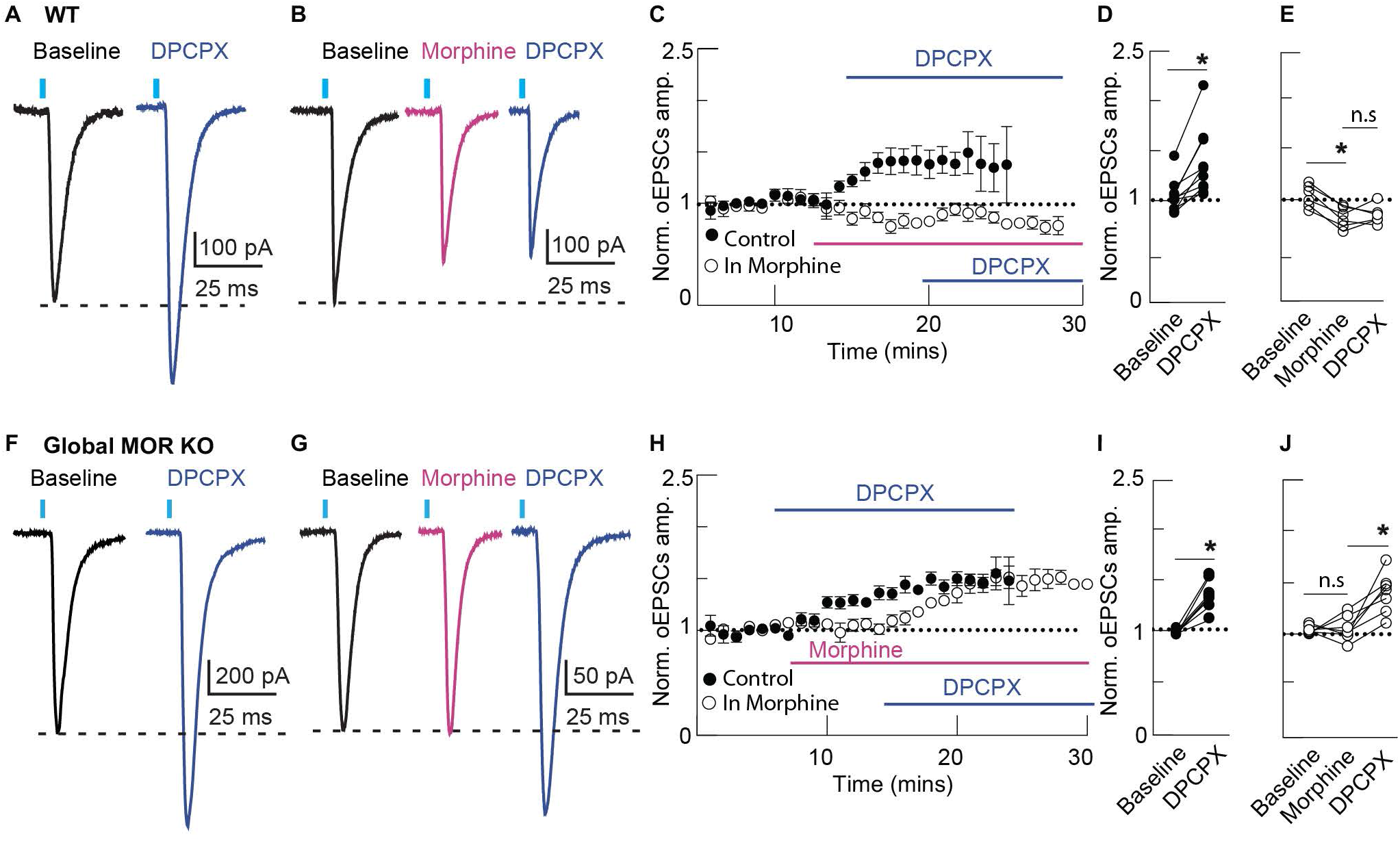
Morphine inhibits adenosine tone in the thalamo-striatal synapse by activating MORs. **(A)** Representative traces of oEPSCs evoked by 470 nm light (black label) and facilitation of oEPSC amplitude by DPCPX (200 nM; blue label). **(B)** Representative traces of oEPSCs evoked by 470 nm light (black label), inhibition of oEPSC amplitude by morphine (1 μM; pink label), and lack of facilitation by DPCPX (200 nM; blue label). **(C)** Plot of the time course of normalized oEPSC amplitude for cells superperfused with DPCPX (dark circles; n = 10 cells, 7 mice), and for cells superperfused with morphine and then DPCPX (clear circle; n = 6 cells, 4mice). **(D)** Mean summary data of normalized oEPSC amplitude in control and after DPCPX (1.3 ± 0.06 fraction of baseline, p = 0.0003, n = 10 cells, 7 mice, t(9) = 5.752, ratio paired T-test). **(E)** Mean summary data of normalized oEPSC amplitude in control, after morphine superperfusion, and after DPCPX superperfusion. Morphine significantly inhibited oEPSC amplitude (morphine 0.78 ± 0.03 fraction of baseline, p = 0.0011; DPCPX: 0.77 ± 0.05 fraction of baseline p = 0.04, and 0.99 ± 0.06 fraction of morphine, p = 0.84, n = 6 cells, 4 mice, F_(2, 10)_ = 14.00, one-way repeated measures ANOVA, Tukey’s multiple comparisons test). **(F)** Representative traces of oEPSCs evoked by 470 nm light (black label) and facilitation of oEPSC amplitude by DPCPX (200 nM; blue label), in slices from global MOR knock-out mice. **(G)** Representative traces of oEPSCs evoked by 470 nm light (black label), lack of inhibition by morphine (1 μM; pink label), and facilitation of oEPSC amplitude by DPCPX (200 nM; blue label), in slices from global MOR knock-out mice. **(H)** Plot of the time course of normalized oEPSC amplitude for cells superperfused with DPCPX (dark circles; n = 8 cells, 5 mice), and for cells superperfused with morphine and then DPCPX (clear circle; n = 6 cells, 3 mice). **(I)** Mean summary data of normalized oEPSC amplitude in control and after DPCPX (1.37 ± 0.05 fraction of baseline, p = 0.0001, n = 8 cells, 5 mice, t(7) = 8.273, ratio paired T-test). **(J)** Mean summary data of normalized oEPSC amplitude in control, after morphine superperfusion, and after DPCPX superperfusion. Morphine did not inhibit oEPSC amplitude (morphine: 1.0 ± 0.05 fraction of baseline, p = 0.9863) and there was facilitation by DPCPX in the presence of morphine (1.4 ± 0.07 fraction of baseline, p = 0.0006, 1.3 ± 0.09 fraction morphine, p = 0.0008, n = 6 cells, 3 mice, F_(2, 12)_ = 17.46, one-way repeated measures ANOVA, Tukey’s multiple comparisons test). Line and error bars represent mean ± SEM; ^*^ denotes statistical significance; ns denotes not significant.

### Tonic endogenous activation of A_1_Rs is regulated by cAMP levels

MOR is a G_i/o_-coupled GPCR that can inhibit adenylyl cyclase so it is possible that morphine decreases tonic adenosine levels by preventing cAMP production and its subsequent metabolism to adenosine. Therefore, the role of cAMP metabolism on A_1_R-mediated inhibition of glutamate release was examined. Slices were pretreated with phosphodiesterase (PDE) inhibitor, R0-20-1724 (400 μM) for at least an hour to block metabolism of cAMP. R0-20-1724 (400 μM) was also in the perfusate throughout the course of the experiment. In the presence of R0-20-1724, unlike in control slices, DPCPX (200 nM) did not cause an increase in oEPSC amplitude (Fig 3A, C, D; DPCPX 1.47 ± 0.13, p = 0.03, n = 6 cells, 4 mice, in control; Fig 3B, C, E; DPCPX 0.96 ± 0.1 fraction of baseline, p = 0.789, n = 6 cells, 4 mice, in R0-201724, F_(3,17)_ = 13.51, one-way repeated measures ANOVA, Tukey’s multiple comparisons test), suggesting that inhibiting the metabolism of cAMP, and thus the conversion of cAMP to adenosine, blocked the tonic activation of A_1_Rs. Exogenous application of adenosine (100 μM) in the either the presence or the absence of R0-20-1724 decreased the oEPSC amplitude, which was reversed by a washout (Fig 3A, B, C, D, E; adenosine 0.52 ± 0.08 fraction of baseline, p = 0.0001; washout: 0.87 ± 0.06 of baseline, p = 0.0001, n = 6 cells, 4 mice in R0-201724; adenosine 0.44 fraction ± 0.07 of baseline, p = 0.0094; washout: 1.0 ± 0.05 of baseline, p = 0.999, n = 6 cells, 4 mice, in control, F_(3, 16)_ = 36.72, one-way repeated measures ANOVA, Tukey’s multiple comparisons test), demonstrating that R0-20-1724 is not directly antagonizing the ability of adenosine to inhibit glutamate release via A_1_Rs.

**Fig 3.**
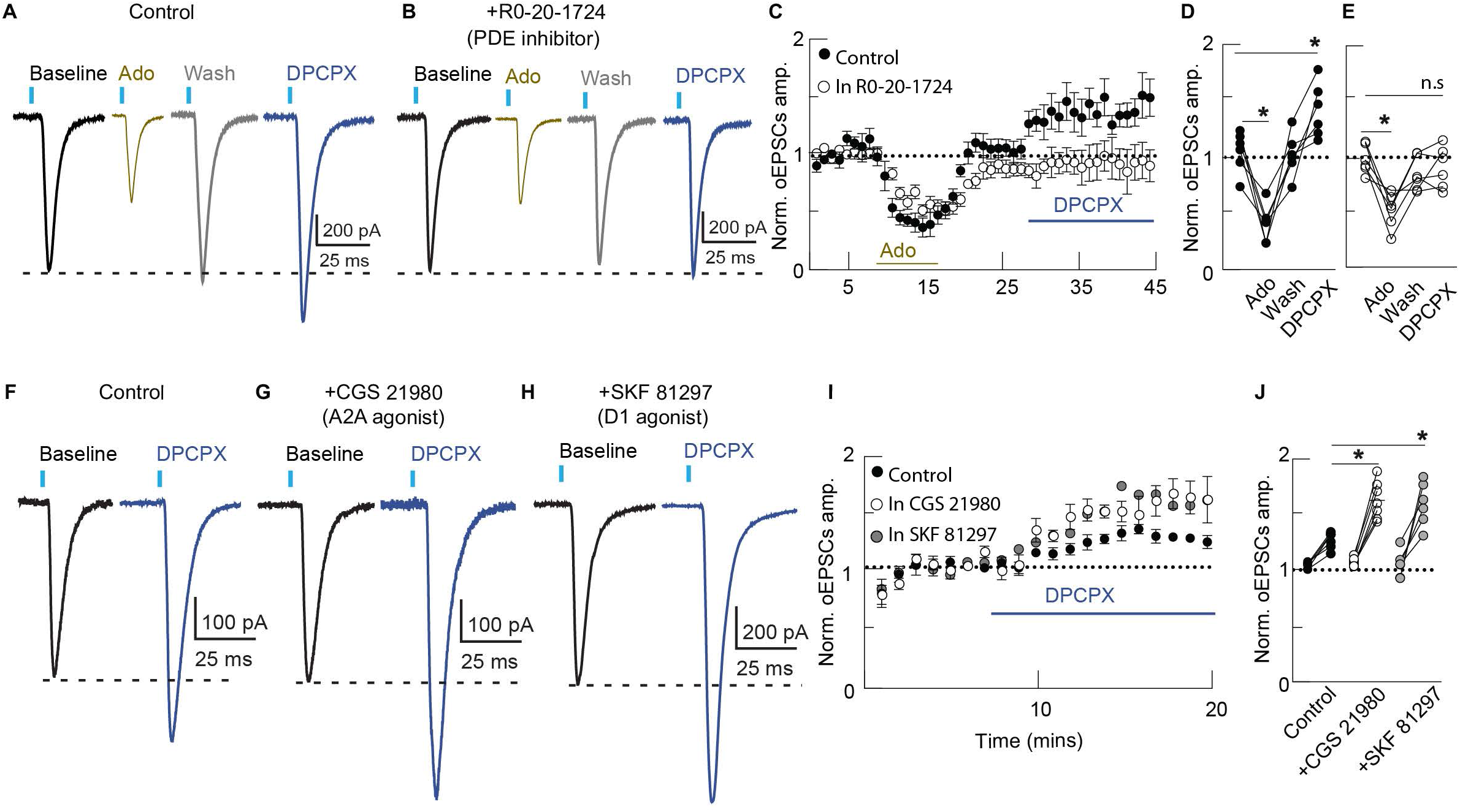
Morphine inhibits adenosine signaling via a cAMP dependent mechanism. **(A)** Representative traces of oEPSCs evoked by 470 nm light (black label), inhibition of oEPSC amplitude by adenosine (100 μM; yellow label), washout of adenosine (gray label), and facilitation of oEPSC by DPCPX (200 nM; blue label) in naïve conditions. **(B)** Representative traces of oEPSCs evoked by 470 nm light (black label), inhibition of oEPSC amplitude by adenosine (100 μ M; yellow label), washout of adenosine (gray label), and lack of facilitation of oEPSC by DPCPX (200 nM; blue label) in slices preincubated in R0-20-1724. **(C)** Plot of the time course of normalized oEPSC amplitude for cells superperfused with adenosine followed by washout and then DPCPX in naïve slices (dark circles, n = 6 cells, 4 Mice) and in slices preincubated in R0-20-1724 (n = 6 cells, 4 mice). **(D)** Mean summary data of normalized oEPSC amplitude for naïve slices in baseline condition, after adenosine superperfusion, followed by a washout and then DPCPX. Adenosine significantly reduced oEPSC amplitude in naïve slices and DPCPX significantly facilitated oEPSC in these slices (adenosine 0.44 fraction ± 0.07 of baseline, p = 0.0094; washout: 1.0 ± 0.05 of baseline, p = 0.999, n = 6 cells, 4 mice, F_(3,17)_ = 13.51, repeated measures ANOVA, Tukey’s multiple comparisons test). **(E)** Mean summary data of normalized oEPSC amplitude for slices incubated in R0-20-1724 in baseline condition, after adenosine superperfusion, followed by a washout and then DPCPX. Adenosine significantly reduced oEPSC amplitude in these slices (adenosine 0.52 ± 0.08 fraction of baseline, p = 0.0001; washout: 0.87 ± 0.06 of baseline, p = 0.0001, n = 6 cells, 4 mice, F_(3,17)_ = 13.51, repeated measures ANOVA, Tukey’s multiple comparisons test), but DPCPX did not significantly facilitate oEPSC in these slices (DPCPX: 0.96 ± 0.10 fraction of baseline, p = 0.8755, repeated measures ANOVA, Tukey’s multiple comparisons test). **(F)** Representative traces of oEPSCs evoked by 470 nm light (black label) and facilitation of oEPSC amplitude by DPCPX (200 nM; blue label), in control slices. **(G)** Representative traces of oEPSCs evoked by 470 nm light (black label) and facilitation of oEPSC amplitude by DPCPX (200 nM; blue label), in slices preincubated in CGS21980. **(H)** Representative traces of oEPSCs evoked by 470 nm light (black label) and facilitation of oEPSC amplitude by DPCPX (200 nM; blue label), in slices preincubated in SKF81290. **(I)** Plot of the time course of normalized oEPSC amplitude for cells superperfused with DPCPX in control condition (dark circles; n = 7 cells, 5 mice), for cells preincubated in SKF81290 (clear circles; n = 7 cells, 6 mice) and for cells preincubated in CGS21980 (gray circles; n = 6 cells, 4 mice). **(J)** Mean summary data of normalized oEPSC amplitude in control, in slices preincubated in SKF81297, and CGS21980. The increase in amplitude induced by DPCPX was significantly higher in slices treated with SKF89217 (DPCPX 1.6 ± 0.11 fraction of baseline, p = 0.003,) and in slices treated with CGS21680 (p < 0.001, F_(5, 32)_ = 32.24, one-way ANOVA, Tukey’s multiple comparisons test) compared to control slices. Line and error bars represent mean ± SEM; ^*^ denotes statistical significance.

In order to examine if endogenous adenosine levels could be increased by increasing cAMP concentration, G_s_ coupled GPCRs in both D_1_R- and D_2_R-positive MSNs were pharmacologically activated. Slices were preincubated in D_1_R specific agonist SKF89217 (1 μM) for at least an hour, with the drug in the perfusate throughout the course of the experiment. DPCPX (200 nM) caused an increase in oEPSC amplitude (Fig 3G, H, I; DPCPX 1.6 ± 0.11 fraction of baseline, n = 7 cells, 5 mice). The increase in amplitude induced by DPCPX was significantly higher in slices treated with SKF89217 (p = 0.003, F_(5, 32)_ = 32.24, one-way ANOVA, Tukey’s multiple comparisons test) compared to control slices. Next, slices were incubated in A_2A_R agonist, CGS21680 (1 μM) for at least an hour, with the drug in the perfusate throughout the course of the experiment. A_2A_Rs co-localize with D_2_R positive MSNs only (Bogenpohl et al. 2012; Fink et al. 1992; Severino et al. 2020). DPCPX (200 nM) increased oEPSC amplitude (Fig 3H, I, J; DPCPX 1.76 ± 0.10 fraction of baseline, n = 6 cells, 4 mice). The increase in amplitude induced by DPCPX was also significantly higher in slices treated with CGS21680 (p < 0.0001, F_(5, 32)_ = 32.24, one-way ANOVA, Tukey’s multiple comparisons test) compared to control slices. There was no difference in oEPSC amplitude after DPCPX superperfusion between slices incubated in SKF89217 and CGS21680 (one-way ANOVA, Tukey’s multiple comparisons test). Thus, basal endogenous adenosine levels are affected by cAMP concentration, and activation of MORs by morphine appears to inhibit cAMP accumulation, consequently decreasing adenosine levels.

### Inhibition of cAMP by activation of MORs is reversible

The time-dependence of cAMP inhibition by MOR activation was examined next. [Met^5^] enkephalin (ME; 1 μM) was used instead of morphine, as ME washes from brain slices. oEPSCs were induced as previously described and ME (1 μM) was superperfused. Like morphine, ME inhibited oEPSCs and DPCPX failed to facilitate oEPSCs in the presence of ME (Fig 4A, B, C; ME 0.67 ± 0.03 fraction of baseline, p = 0.0002; DPCPX 0.56 ± 0.03 fraction of baseline, p = < 0.0001; DPCPX 0.84 ± 0.05 fraction of ME, p = 0.2867, n = 6 cells, 4 mice, F_(3, 15)_ = 78.77, repeated measures one-way ANOVA, Tukey’s multiple comparisons test). ME washed out of the slices in approximately five minutes. Following ME washout, there was an over-reversal of oEPSC in the presence of DPCPX (DPCPX 1.37 ± 0.07 fraction of baseline, p = <0.0001, repeated measures one-way ANOVA, Tukey’s multiple comparisons test) which reached steady-state approximately in seven minutes, suggesting that inhibition of cAMP, and therefore inhibition of tonic adenosine levels, is acutely reversible.

**Fig 4.**
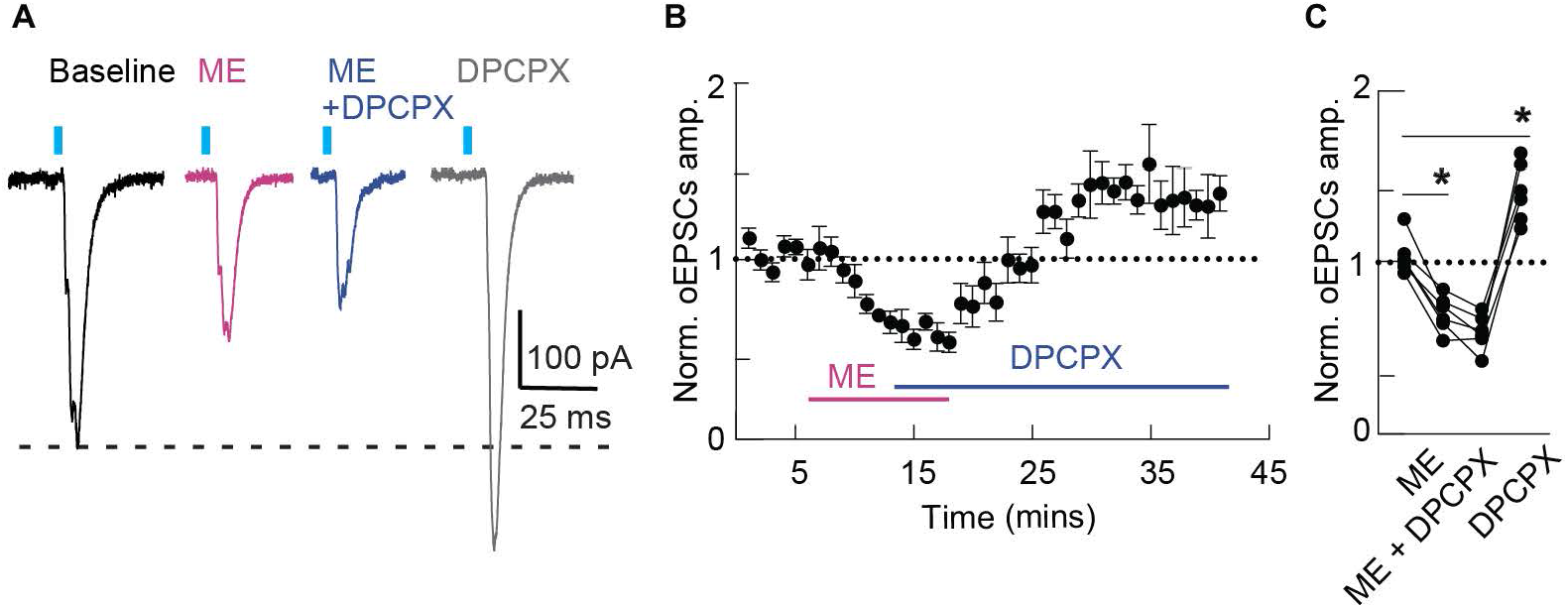
Inhibition of adenosine signaling by opioids is reversible. **(A)** Representative traces of oEPSCs evoked by 470 nm light (black label), inhibition of oEPSC amplitude by ME (1 μM; pink label), lack of facilitation by DPCPX (200 nM; blue label), and an over-reversal of oEPSC after ME washout (gray label). **(B)** Plot of the time course of normalized oEPSC amplitude for cells superperfused with ME, followed by DPCPX in the presence of ME, and then a washout of ME, but not DPCPX (n = 6 cells, 4 mice). **(C)** Mean summary data of normalized oEPSC amplitude in baseline condition, after ME superperfusion, followed by DPCPX, and a washout of ME, but not DPCPX (ME 0.67 ± 0.03 fraction of baseline, p = 0.0002; DPCPX 0.56 ± 0.03 fraction of baseline, p = < 0.0001; DPCPX 0.84 ± 0.05 fraction of ME, p = 0.2867, n = 6 cells, 4 mice, F_(3, 15)_ = 78.77, repeated measures one-way ANOVA, Tukey’s multiple comparisons test). Line and error bars represent mean ± SEM; ^*^ denotes statistical significance; ns denotes not significant.

### Delta and kappa opioid receptors do not regulate tonic activation of adenosine A_1_ receptors

Next, the effect on tonic activation of A_1_Rs by delta opioid receptor (DOR) and kappa opioid receptor (KOR) activation were examined. oEPSCs were evoked as described above and the DOR selective agonist deltorphin (300 nM) was superperfused, followed by DPCPX (200 nM). Unlike morphine, deltorphin did not reduce the amplitude of oEPSCs and, in the presence of deltorphin, DPCPX increased oEPSC amplitude (Fig 5A, C, D; deltorphin 1.0 ± 0.04 fraction of baseline, p = 0.90; DPCPX: 1.42 ± 0.07 fraction of baseline, p = 0.0002, and 1.40 ± fraction of deltorphin, p = 0.0004, n = 6 cells, 3 mice, F_(2, 10)_ = 24.60, repeated measures one-way ANOVA, Tukey’s multiple comparisons test), suggesting that DOR activation does not affect the tonic activation of A_1_Rs. Next, in separate cells, the KOR selective agonist U69,593 (1 μM) was superperfused, followed by DPCPX (200 nM). Similar to deltorphin, U69,593 did not inhibit oEPSC, and in the presence of U69,593 DPCPX increased oEPSC amplitude (Fig 5B, C, E; U69,593 1.02 ± 0.06 fraction of baseline p = 0.9670; DPCPX: 1.6 ± 0.09 fraction of baseline, p = 0.0051, and 1.6 ± 0.13 fraction of U69,593, p = 0.0035, n = 6 cells, 4 mice, F_(2, 10)_ = 12.24, repeated measures one-way ANOVA, Tukey’s multiple comparisons test), suggesting that KOR activation, like DOR activation, did not inhibit glutamate release from thalamic terminals or affect the tonic activation of A_1_Rs. Hence, both the direct inhibition of glutamate release from thalamic afferents and the inhibition of tonic activation of A_1_Rs seems to be agonist specific, both inhibited only by MOR agonists and not DOR or KOR agonists.

**Fig 5.**
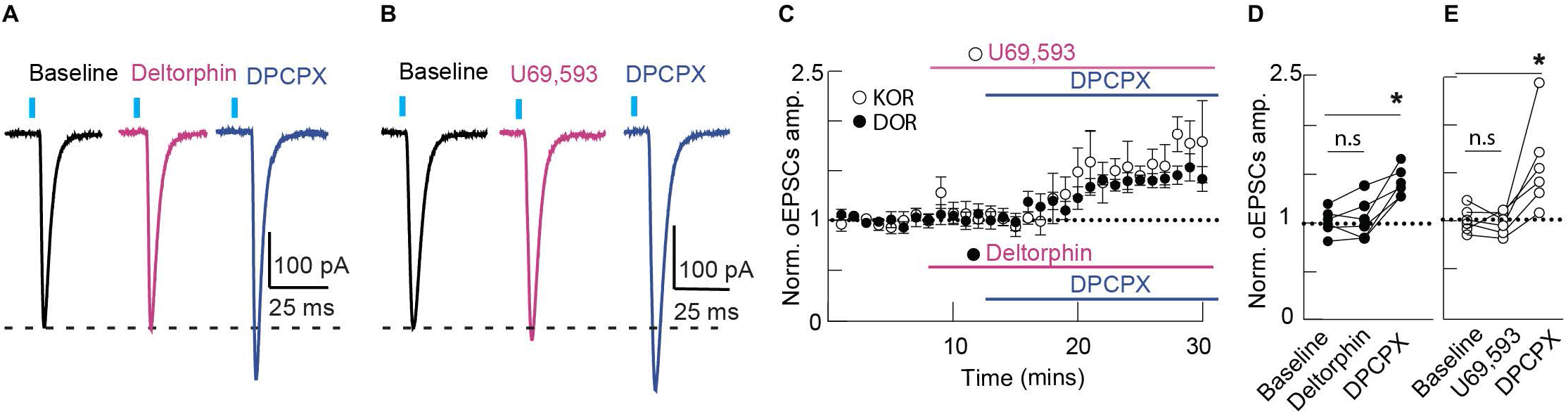
DORs and KORs do not mediate inhibition of adenosine signaling. **(A)** Representative traces of oEPSCs evoked by 470 nm light (black label), lack of inhibition of oEPSC amplitude by deltorphin (300 nM; pink label), and facilitation by DPCPX (200 nM; blue label). **(B)** Representative traces of oEPSCs evoked by 470 nm light (black label), lack of inhibition of oEPSC amplitude by U69,593 (1 μM; pink label), and facilitation by DPCPX (200 nM; blue label). **(C)** Plot of the time course of normalized oEPSC amplitude for cells superperfused with deltorphin (black circles), followed by DPCPX (n = 6 cells, 3 Mice), and for cells superperfused with U69 (clear circles), followed by DPCPX (n = 6 cells, 4 Mice). **(D)** Mean summary data of normalized oEPSC amplitude in baseline condition, after deltorphin superperfusion, followed by DPCPX (deltorphin 1.0 ± 0.04 fraction of baseline, p = 0.90; DPCPX: 1.42 ± 0.07 fraction of baseline, p = 0.0002, and 1.40 ± fraction of deltorphin, p = 0.0004, n = 6 cells, 3 mice, F_(2, 10)_ = 24.60, repeated measures one-way ANOVA, Tukey’s multiple comparisons test). **(E)** Mean summary data of normalized oEPSC amplitude in baseline condition, after U69,593 superperfusion, followed by DPCPX (U69,593 1.02 ± 0.06 fraction of baseline p = 0.9670; DPCPX: 1.6 ± 0.09 fraction of baseline, p = 0.0051, and 1.6 ± 0.13 fraction of U69,593, p = 0.0035, n = 6 cells, 4 mice, F_(2, 10)_ = 12.24, repeated measures one-way ANOVA, Tukey’s multiple comparisons test). Line and error bars represent mean ± SEM; ^*^ denotes statistical significance; ns denotes not significant.

### Presynaptic effects of MOR agonists in the thalamo-striatal circuit

Because the inhibition of tonic adenosine release by opioids was selectively mediated by MORs, and since these receptors are expressed in the thalamic terminals, and both the D_1_R-positive and D_2_R-positive MSNs, the location of acute action of MOR agonist was investigated. FloxedMOR (*Oprm1*^*fl/fl*^) mice and Vglut_2_:*cre* mice were crossed to generate mice lacking MORs from Vglut_2_-expressing presynaptic terminals (*Oprm1*^*fl/fl*^, Vglut_2_-*cre* ^*+/-*^) (Vong et al. 2011). FloxedMOR homozygous littermates that did not express Vglut_2_:*cre* were used as controls (*Oprm1*^*fl/fl*^, Vglut_2_-*cre* ^*-/-*^). oEPSCs were evoked as previously described. Superperfusion of the MOR agonist DAMGO (1 μM) decreased the amplitude of the oEPSCs, and this inhibition was reversed by the antagonist naloxone (1 μM) (Fig 6A, C, D; DAMGO 0.39 ± 0.05 fraction of baseline, p < 0.0001; naloxone: 0.80 ± 0.06 of baseline, n = 8 cells, 4 mice, F_(2, 14)_ = 29.9, repeated measures one-way ANOVA, Tukey’s multiple comparisons test), in control mice. In FloxedMOR-Vglut_2_-*cre* mice lacking MORs in the presynaptic terminals, DAMGO did not inhibit the amplitude of the oEPSCs (Fig 4B, C, E; DAMGO: 0.99 ± 0.02 fraction of baseline, p = 0.6023; naloxone: 0.92 ± 0.05 fraction of baseline, p = 0.14, n = 7 cells, 6 mice, F_(2, 12)_ = 2.08, repeated measures one-way ANOVA, Tukey’s multiple comparisons test), suggesting that opioid action on the thalamo-striatal glutamate release is presynaptic and demonstrating the effectiveness of cre-dependent knockout in these animals. Next, in order to examine if adenosine was released from presynaptic terminals, DPCPX (200 nM) was superperfused. DPCPX increased the amplitude of the oEPSCs in the presynaptic MOR KO mice (Fig 6F, H, I; DPCPX: 1.3 ± 0.07 fraction of baseline, p = 0.0012, n = 8 cells, 4 mice, t(7) = 5.225, ratio paired T-test). In separate cells, morphine (1 μM) was superperfused, followed by DPCPX. As expected, morphine did not reduce the amplitude of oEPSCs, however, in the presence of morphine, DPCPX did not increase the amplitude of oEPSCs (Fig 6G, H, J; morphine 1.0 ± 0.07 fraction of baseline, p = 0.9935; DPCPX: 1.0 ± 0.07 fraction of baseline, p = 0.9119, and 1.0 ± 0.09 fraction of morphine, p = 0.9513, n = 6 cells, 4 mice, F_(2, 10)_ = 0.09141, repeated measures one-way ANOVA, Tukey’s multiple comparisons test), suggesting that morphine still inhibited the tonic activation of A_1_Rs, even in mice lacking presynaptic MORs. Therefore, even though opioids presynaptically inhibit glutamate release from the thalamic terminals, the presynaptic MORs do not regulate extracellular adenosine accumulation, and subsequent tonic activation of the A_1_Rs in this circuit.

**Figure 6.**
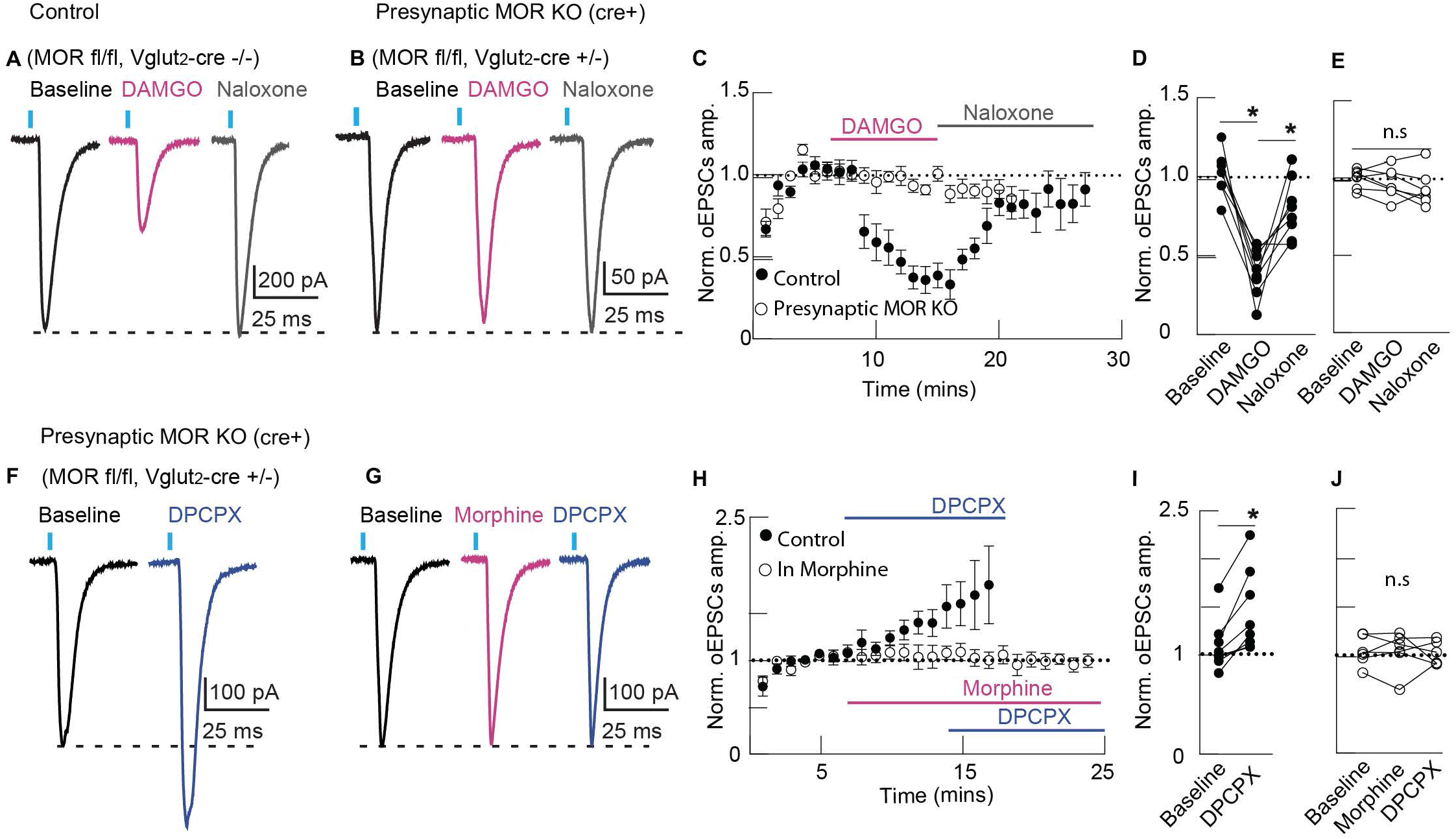
Presynaptic MORs suppress excitatory thalamic inputs, but do not regulate tonic A_1_R activation. **(A)** Representative traces of oEPSCs evoked by 470 nm light (black label), inhibition of oEPSC amplitude by DAMGO (1 μM; pink label), and reversal by naloxone (1 M; gray label) in control mice expressing MORs in presynaptic terminals. **(B)** Representative traces of oEPSCs evoked by 470 nm light (black label), lack of inhibition of oEPSC amplitude by DAMGO (1 μ M; pink label), and no effect of naloxone (1 M; gray label) in mice lacking MORs in presynaptic terminals. **(C)** Plot of the time course of normalized oEPSC amplitude for cells superperfused with DAMGO followed by naloxone in control mice (dark circles, n = 8 cells, 4 mice) and in mice lacking MORs in presynaptic terminals (clear circles, n = 7 cells, 6 mice). **(D)** Mean summary data of normalized oEPSC amplitude in for control mice in baseline condition, after DAMGO superperfusion, followed by naloxone (DAMGO 0.39 ± 0.05 fraction of baseline, p < 0.0001; naloxone: 0.80 ± 0.06 of baseline, n = 8 cells, 4 mice, F_(2, 14)_ = 29.9, repeated measures one-way ANOVA, Tukey’s multiple comparisons test). **(E)** Mean summary data of normalized oEPSC amplitude for presynaptic MOR KO mice in baseline condition, after DAMGO perfusion, followed by Naloxone (DAMGO: 0.99 ± 0.02 fraction of baseline, p = 0.6023; naloxone: 0.92 ± 0.05 fraction of baseline, p = 0.14, n = 7 cells, 6 mice, F_(2, 12)_ = 2.08, repeated measures one-way ANOVA, Tukey’s multiple comparisons test). Line and error bars represent mean ± SEM; * denotes statistical significance; ns denotes not significant. **(F)** Representative traces of oEPSCs evoked by 470 nm light (black label) and facilitation of oEPSC amplitude by DPCPX (200 nM; blue label) in mice lacking MORs in presynaptic terminals. **(G)** Representative traces of oEPSCs evoked by 470 nm light (black label), lack of inhibition of oEPSC amplitude by morphine (1 μM; pink label), and lack of facilitation by DPCPX (200 nM; blue label). **(H)** Plot of the time course of normalized oEPSC amplitude for cells superperfused with DPCPX (dark circles; n = 8 cells, 4 mice), and for cells superperfused with morphine and then DPCPX (clear circle; n = 6 cells, 4 mice). **(I)** Mean summary data of normalized oEPSC amplitude in control and after DPCPX (1.3 ± 0.07 fraction of baseline, p = 0.0012, n = 8 cells, 4 mice, t(7) = 5.225, ratio paired T-test). **(J)** Mean summary data of normalized oEPSC amplitude in control, after morphine superperfusion, followed by DPCPX. Morphine did not inhibit oEPSC amplitude (morphine 1.0 ± 0.07 fraction of baseline, p = 0.9935), and there was no facilitation by DPCPX in the presence of morphine (1.0 ± 0.07 fraction of baseline, p = 0.9119, and 1.0 ± 0.09 fraction of morphine, p = 0.9513, n = 6 cells, 4 mice, F_(2, 10)_ = 0.09141, repeated measures one-way ANOVA, Tukey’s multiple comparisons test). Line and error bars represent mean ± SEM; ^*^ denotes statistical significance; ns denotes not significant.

### μ opioid receptor sensitive adenosine release is regulated by D_1_ receptor-expressing MSNs, not D_2_ receptor-expressing MSNs

MORs are expressed in both D_1_ and D_2_ receptor expressing MSNs (Cui et al. 2014; Oude Ophuis et al. 2014), and activation of D_1_ and A_2A_ receptors, presumably in D_1_ and D_2_ receptor expressing MSNs, increased tonic adenosine inhibition of A_1_Rs (Fig 3B), suggesting that MSNs are the potential source of extracellular adenosine. Therefore, MORs were selectively knocked-out in these cells. FloxedMOR mice and A_2A_:*cre* mice were crossed to generate mice lacking MORs from D_2_R expressing MSNs (*Oprm1*^*fl/fl*^, A_2A_-*cre* ^*+/-*^) (Gong et al. 2007). oEPSCs were evoked as previously described, and DPCPX increased the amplitude of the oEPSCs (Fig 7A, C, D; DPCPX: 1.3 ± 0.05 fraction of baseline, p = 0.0049, n = 6 cells, 4 mice, t(5) = 4.787, ratio paired T-test). In separate cells, morphine (1 μM) was superperfused, followed DPCPX. Morphine reduced the amplitude of oEPSCs and, in the presence of morphine, DPCPX did not increase the amplitude of oEPSCs (Fig 7B, C, F; morphine 0.76 ± 0.03 fraction of baseline, p = 0.0001; DPCPX: 0.75 ± 0.02 fraction of baseline, and 0.99 ± 0.04 fraction of morphine, p = 0.9969, n = 6 cells, 4 mice, F_(2, 10)_ = 30.38, repeated measures one-way ANOVA, Tukey’s multiple comparisons test), suggesting that morphine inhibited the tonic activation of A_1_Rs in mice lacking MORs in D_2_R-expressing MSNs. Next, FloxedMOR mice and D_1_: *cre* mice were crossed to generate mice lacking MORs from D_1_R expressing MSNs (*Oprm1*^*fl/fl*^, D_1_-*cre*^*+/-*^) (Gong et al. 2007). DPCPX (200 nM) increased the amplitude of the oEPSCs (Fig 7F, H, I; DPCPX; 1.4 ± 0.09 fraction of baseline, p = 0.006, n = 5 cells, 3 mice, t(4) = 5.253, ratio paired T-test). In sperate cells, morphine (1 μM) reduced the amplitude of oEPSCs and, in the presence of morphine, unlike in WT mice, DPCPX increased the amplitude of oEPSCs (Fig 7G, H, J; morphine 0.72 ± 0.04 fraction of baseline, p = 0.013; DPCPX: 1.14 ± 0.06 fraction of baseline, p = 0.06 and 1.51 ± 0.08 fraction of morphine, p = 0.0001, 5 cells, 3 mice, F_(3, 12)_ = 25.36, repeated measures one-way ANOVA, Tukey’s multiple comparisons test). Next, MOR antagonist naloxone caused an over-reversal of oEPSC compared to baseline (Fig 7G, H, J; naloxone 1.30 ± 0.03 fraction of baseline, p = 0.004), suggesting that in mice lacking MOR in D_1_R positive MSNs, morphine could no longer inhibit tonic A_1R_ activation. Surprisingly, mice lacking MORs in only one copy of the D_1_R gene (FloxedMOR +/-, D_1_-cre +/-), also showed similar results. In these mice, DPCPX (200 nM) increased the amplitude of the oEPSCs as well (Fig 7F, H, I; DPCPX; 1.38 ± .22 fraction of baseline, p < .001, n = 6 cells, 5 mice, t(5) = 4.466, ratio paired T-test). In sperate cells, morphine (1 μM) reduced the amplitude of oEPSCs and, in the presence of morphine, unlike in WT mice, DPCPX increased the amplitude of oEPSCs (Fig 7G, H, J; morphine .70 ± .05 fraction of baseline, p = .0003; DPCPX: .91 ± .06 fraction of baseline, p = .39, and 1.34 ± .06 fraction of morphine, p = .0086, 6 cells, 4 mice, F_(3, 15)_ = 27.12, repeated measures one-way ANOVA, Tukey’s multiple comparisons test). Next, MOR antagonist naloxone caused an over-reversal of oEPSC compared to baseline (Fig 7G, H, J; naloxone 1.2 ± .05 fraction of baseline, p = 0.0224), suggesting that a partial deletion of MORs from D_1_R expressing MSNs was sufficient to eliminate the inhibition of tonic A_1_R signaling. Combined, these results demonstrate that morphine-mediated adenosine release in the thalamo-striatal circuit comes from D_1_R positive MSNs.

**Figure 7.**
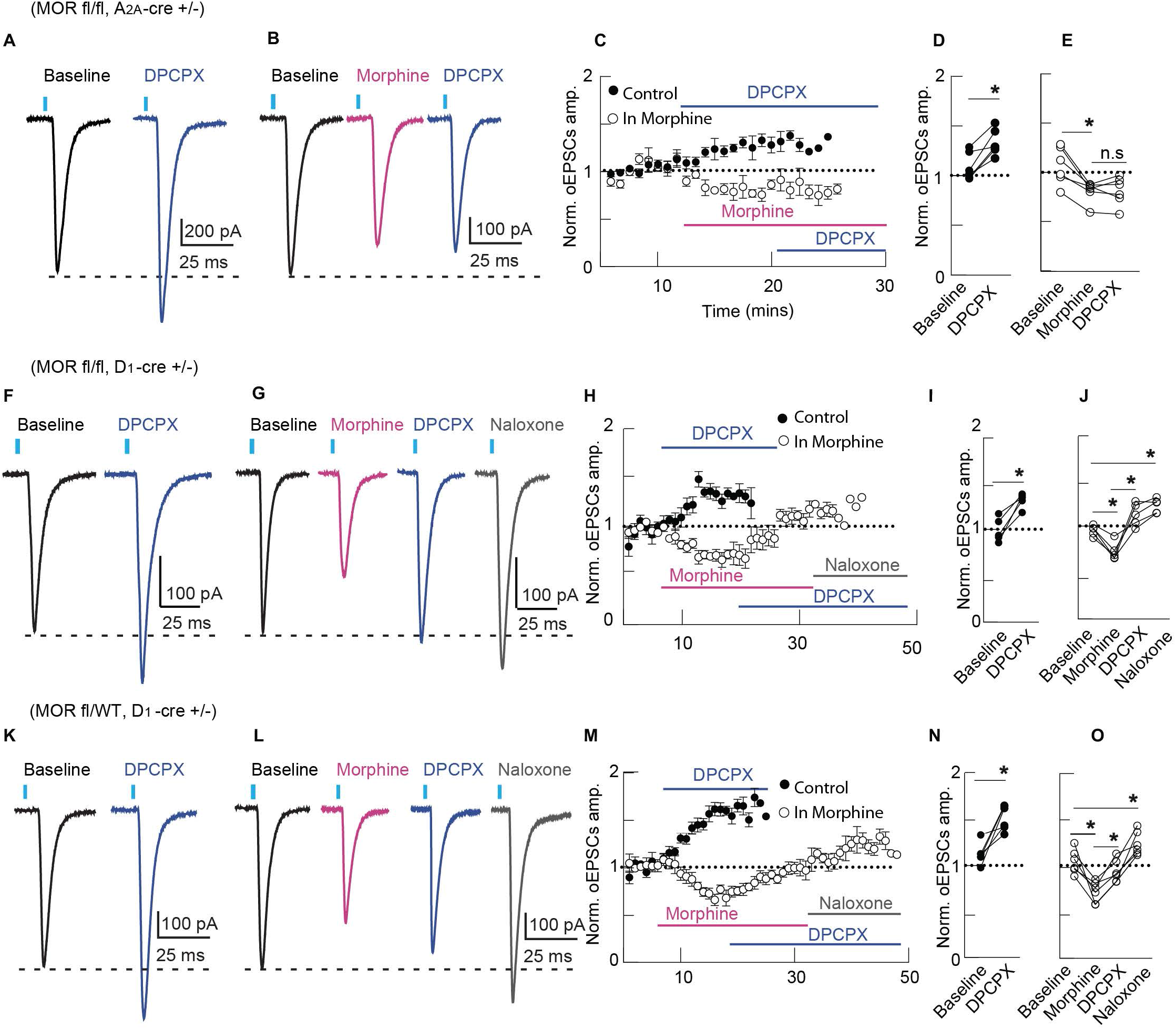
MORs in D_1_R expressing MSNs, but not D_2_R expressing MSNs, regulate tonic A_1_R activation. **(A)** Representative traces of oEPSCs evoked by 470 nm light (black label), facilitation of oEPSC amplitude by DPCPX (100 nM; blue label mice lacking MORs in D_2_R expressing MSNs. **(B)** Representative traces of oEPSCs evoked by 470 nm light (black label), inhibition of oEPSC amplitude by morphine (1 μM; pink label), and lack of facilitation by DPCPX (200 nM; blue label) in mice lacking MORs in D_2_R expressing MSNs. **(C)** Plot of the time course of normalized oEPSC amplitude for cells superperfused with DPCPX (dark circles; n = 6 cells, 4 mice), and for cells superperfused with morphine and then DPCPX (clear circle; n = 6 cells, 4 mice). **(D)** Mean summary data of normalized oEPSC amplitude in control and after DPCPX (DPCPX: ± 0.05 fraction of baseline, p = 0.0049, n = 6 cells, 4 mice, t(5) = 4.787, ratio paired T-test). **(E)** Mean summary data of normalized oEPSC amplitude in control, after morphine superperfusion, and after DPCPX superperfusion. Morphine significantly inhibited oEPSC amplitude (morphine 0.76 ± 0.03 fraction of baseline, p = 0.0001), but there was no facilitation by DPCPX in the presence of morphine (DPCPX: 0.75 ± 0.02 fraction of baseline, and 0.99 ± 0.04 fraction of morphine, p = 0.9969, n = 6 cells, 4 mice, F_(2,10)_ = 30.38, repeated measures one-way ANOVA, Tukey’s multiple comparisons test). **(F)** Representative traces of oEPSCs evoked by 470 nm light (black label) and facilitation of oEPSC amplitude by DPCPX (200 nM; blue label), in slices from mice lacking MORs from D_1_R expressing MSNs. **(G)** Representative traces of oEPSCs evoked by 470 nm light (black label), inhibition by morphine (1 μ M; pink label), and facilitation of oEPSC amplitude by DPCPX (200 nM; blue label), in slices from mice lacking MORs from D_1_R expressing MSNs. **(H)** Plot of the time course of normalized oEPSC amplitude for cells superperfused with DPCPX (dark circles; n = 5 cells, 3 mice), and for cells superperfused with morphine and then DPCPX (clear circle; n = 5 cells, 3 mice). **(I)** Mean summary data of normalized oEPSC amplitude in control and after DPCPX (DPCPX; ± 0.09 fraction of baseline, p = 0.006, n = 5 cells, 3 mice, t(4) = 5.253, ratio paired T-test). **(J)** Mean summary data of normalized oEPSC amplitude in control, after morphine superperfusion, and after DPCPX superperfusion. Morphine inhibited oEPSC amplitude (morphine 0.72 ± 0.04 fraction of baseline, p = 0.013;) and there was facilitation by DPCPX in the presence of morphine (1.14 ± 0.06 fraction of baseline, p = 0.06 and 1.51 ± 0.08 fraction of morphine, p = 0.0001, 5 cells, 3 mice, F_(3, 12)_ = 25.36, repeated measures one-way ANOVA, Tukey’s multiple comparisons test). Naloxone caused an over-reversal of oEPSC amplitude (1.30 ± 0.03 fraction of baseline, p = 0.004). Line and error bars represent mean ± SEM; * denotes statistical significance; ns denotes not significant. **(K)** Representative traces of oEPSCs evoked by 470 nm light (black label) and facilitation of oEPSC amplitude by DPCPX (200 nM; blue label), in slices from mice with a partial MOR knock-out from D_1_R expressing MSNs. **(L)** Representative traces of oEPSCs evoked by 470 nm light (black label), inhibition by morphine (1 μM; pink label), and facilitation of oEPSC amplitude by DPCPX (200 nM; blue label), in slices from mice with a partial MOR knock-down from D_1_R expressing MSNs. **(M)** Plot of the time course of normalized oEPSC amplitude for cells superperfused with DPCPX (dark circles; n = 3 cells, 2 mice), and for cells superperfused with morphine and then DPCPX, followed by naloxone (clear circle; n = 3 cells, 2 mice). **(N)** Mean summary data of normalized oEPSC amplitude in control and after DPCPX (1.38 ± .22 fraction of baseline, p < .001, n = 6 cells, 5 mice, t(5) = 4.466, ratio paired T-test). **(O)** Mean summary data of normalized oEPSC amplitude in control, after morphine superperfusion, after DPCPX superperfusion, and after naloxone superperfusion. Morphine inhibited oEPSC amplitude (morphine .70 ± .05 fraction of baseline, p = .0003) and there was facilitation by DPCPX in the presence of morphine (DPCPX: .91 ± .06 fraction of baseline, p = .39, and 1.34 ± .06 fraction of morphine, p = .0086, 6 cells, 4 mice, F_(3, 15)_ = 27.12, repeated measures one-way ANOVA, Tukey’s multiple comparisons test). Naloxone caused an over-reversal of oEPSC amplitude (1.2 ± .05 fraction of baseline, p = 0.0224). Line and error bars represent mean ± SEM; ^*^ denotes statistical significance; ns denotes not significant.

A summary of the effects of selective deletion of MOR from various neuronal populations demonstrates that while the effect of DPCPX was similar in the absence of morphine across all genotypes, only selective knockout of MOR in D_1_R-positive cells resulted in a significant effect of DPCPX in the presence of morphine compared to WT mice (Fig 8, *Oprm1*^*fl/fl*^, D_1_-*cre*^*+/-*^ p = 0.0004). Additionally, there was no statistical difference between *Oprm1*^*fl/fl*^, D_1_-*cre*^*+/-*^ mice and global MOR KO mice in morphine condition (p = 0.3653, unpaired T-test, t(10) = 0.9485). Combined, these results demonstrate that morphine’s regulation of adenosine signaling in this thalamo-striatal circuit critically requires MORs in D_1_R positive MSNs and that, under these experimental conditions, these D_1_R-positive neurons are the likely source of extracellular adenosine accumulation in dorsomedial striatum.

**Figure 8.**
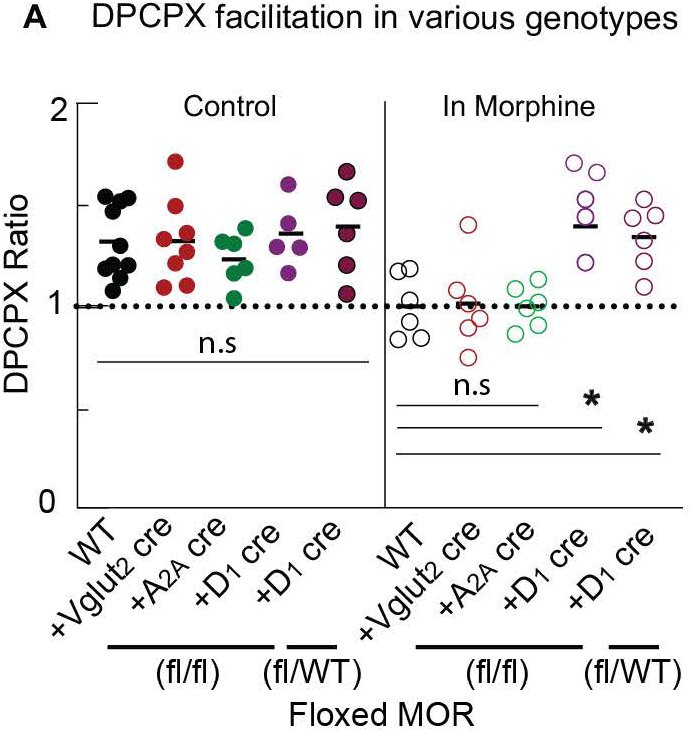
Summary data comparing DPCPX response in the absence and presence of morphine in mice across all genotypes. **(A)** Ratio of DPCPX facilitation compared to baseline and in morphine condition in WT mice, mice lacking MORs in presynaptic thalamic terminals, postsynaptic D_1_R expressing MSNs and D_2_R expressing genotypes. There was no difference in DPCPX responses across genotypes in baseline condition, but mice lacking MORs in D_1_R positive MSNs had a higher facilitation by DPCPX in morphine condition compared to WT mice, mice lacking MORs in presynaptic thalamic terminals and postsynaptic D_2_R expressing MSNs (p = 0.03 compared to WT, p = 0.04 compared to presynaptic MOR KOs, and p = 0.03 compared to MOR KO in D_2_R positive MSNs, one-way ANOVA, Tukey’s multiple comparisons test). Line and error bars represent mean ± SEM; ^*^ denotes statistical significance; ns denotes not significant.

## Discussion

This study explored how the opioid and adenosine signaling systems interact to inhibit glutamate release in a thalamo-striatal circuit. Consistent with previous findings, activation of MORs but not DORs inhibited glutamate release from thalamic terminals (Birdsong et al. 2019). There also was no effect on glutamate release when KOR agonists were superperfused, suggesting the lack of KORs in the thalamo-striatal projections from medial thalamus to dorsomedial striatum. Additionally, activation of A_1_Rs also inhibited glutamate release and antagonism of this receptor revealed endogenous adenosine tone that activated the A_1_Rs. Opioids inhibited this tonic A_1_R activation through MOR, but not DOR or KOR, via a cAMP-mediated mechanism. When MORs were selectively knocked-out from presynaptic terminals and D_2_R positive postsynaptic medium spiny neurons (MSNs), morphine-mediated inhibition of tonic A_1_R activation remained. In contrast, in mice lacking MORs in the D_1_R positive MSNs, morphine no longer inhibited the tonic activation of A_1_Rs. Thus, morphine-sensitive tonic endogenous adenosine in the thalamo-striatal circuit likely arises from D_1_R positive MSNs.

### Interaction between opioids, cAMP, and adenosine

There is evidence for increased basal endogenous adenosine after chronic morphine treatment and withdrawal (Bonci and Williams 1996; Chieng and Williams 1998; Matsui et al. 2014), therefore, acute morphine application having an opposite effect of decreasing cAMP concentration, and subsequently adenosine release, is consistent with the results of this study. However, it should be noted that there is also evidence for cell-type specificity in the striatum after acute and chronic treatment by morphine, with cAMP concentration increasing in the D_1_R positive MSNs in acute morphine condition, and cAMP concentration increasing in the D_2_R positive MSNs in chronic morphine condition (Muntean, Dao, and Martemyanov 2019). Further, MOR activation is known to decrease AC activity and consequently, cAMP accumulation. The role of cAMP as a precursor for extracellular adenosine has been previously established in the hippocampus (Brundege et al. 1997; Brundege and Dunwiddie 1998; Dunwiddie, Diao, and Proctor 1997). Therefore, it is not surprising that fluctuation in cAMP concentration mediates tonic adenosine levels in dorsal medial striatum as well. Previous studies have shown that cAMP metabolism and transport alter adenosine concentration, and that the regulation of extracellular adenosine depends, in part, on the balance between mechanisms that increase and decrease cAMP concentration (Rosenberg and Dichter 1989; Krupinski et al. 1989).

Additionally, in the hippocampus, endogenous adenosine inhibits glutamate release and the basal concentration of endogenous adenosine is about 200 nM (Dunwiddie and Diao 1994). Similarly, there is evidence for basal endogenous adenosine affecting some striatal synapses. For example, there was a potentiation by DPCPX in glutamate release in nucleus accumbens core and GABA release in nucleus accumbens core and shell (Brundege and Williams 2002). This study confirmed that DPCPX also potentiated glutamate release from thalamic terminals in the dorsal medial striatum and a similar cAMP-dependent mechanism mediated adenosine accumulation like in the hippocampus.

### Opioid selectivity in mediating adenosine release in thalamo-striatal circuit

Consistent with previous findings, activation of MOR, but not DOR, led to inhibition of glutamate release in the thalamo-striatal circuit (Birdsong et al. 2019). Furthermore, lack of presynaptic inhibition of glutamate release in FloxedMor-Vglut_2_-*cre* mice corroborates previous finding that MORs in thalamic glutamate terminals regulate transmitter release (Reeves et al. 2020). Additionally, though opioids did not inhibit glutamate release in presynaptic MOR KO mice, morphine still inhibited adenosine tone. Though the thalamic terminals may not express detectable levels of functional DORs, both the D_2_R positive MSNs, and cholinergic interneurons are enriched in DORs (Bertran-Gonzalez et al. 2013), with DORs in the patch region of the striatum inhibiting GABA release from MSN collaterals (Banghart et al. 2015). Additionally, activation of KOR did not inhibit glutamate release, suggesting that effect of opioids on this thalamo-striatal circuit is agonist specific. The dynorphin system in the nucleus accumbens has been implicated in both aversive and rewarding behavior (Al-Hasani et al. 2015), but the circuit and cell-type specificity driving these opposing behaviors is unknown and a potential avenue of future studies. Additionally, neither the activation of DOR nor KOR inhibited tonic adenosine release, suggesting that the MOR uniquely interacts with the adenosine system. The lack of effect of DOR agonists on tonic adenosine release also supports the observation that DORs appear to be enriched in D_2_R expressing rather than D_1_R expressing MSNs (Banghart et al. 2015). Lastly, while the results here have been consistent and reproducible across slices and animals, all experiments were performed in the dorsomedial striatum. It is possible that regional heterogeneity exists in opioid-regulation of adenosine tone such that differences may exist between medial and lateral dorsal striatum or nucleus accumbens.

### Source of opioid-sensitive adenosine tone

Selectively knocking-out the MORs from the presynaptic terminals, D_1_R positive MSNs, and D_2_R positive MSNs revealed that morphine-induced inhibition of adenosine release was due to morphine’s action on MORs in the D_1_R positive MSNs, but not D_2_R positive MSNs. This finding is consistent with previous work showing that D_1_R and D_2_R positive MSNs differentially modulate striatal activity (Lobo and Nestler 2011), and that the somatodendritic region of neurons can release adenosine and retrogradely bind presynaptic A_1_Rs (Lovatt et al. 2012). Furthermore, MORs in D_1_R positive MSNs and D_2_R positive MSNs also differentially modulate opioid responses. MOR deletion from D_1_R positive MSNs inhibits opioid-induced hyperlocomotion while deletion from D_2_R positive MSNs increase opioid-induced hyperlocomotion (Severino et al. 2020). Additionally, MOR expression in the D_1_R positive MSNs was shown to be necessary for opioid self-administration and reward (Cui et al. 2014) Thus, a novel role for MORs in regulating adenosine release in the striatum in a cell-type specific way can have profound implication for opioid dependence and addiction. It is also important to note that there is evidence for astrocytes mediating adenosine release in nucleus accumbens, though the mechanism behind adenosine release is through increases in Ca^2+^ activity, and not through an increase in cAMP concentration (Corkrum et al. 2020). The similarities and differences in ways adenosine is regulated to maintain homeostasis in striatal neuron signaling could be a potential new area of study.

The present results indicate that morphine inhibits tonic adenosine release by activating MORs, and subsequently inhibiting cAMP. This effect of opioid-induced inhibition of adenosine release was specific to MOR and not mediated by DOR or KOR. Selective KO of MORs from presynaptic terminals showed that though opioids presynaptically inhibit glutamate release, presynaptic MORs do not modulate extracellular adenosine accumulation and adenosine signaling in the thalamo-striatal circuit. Rather, tonic adenosine release was no longer inhibited by morphine when MORs were knocked-out from D_1_R positive MSNs, but not D_2_Rs positive MSNs or from glutamate terminals. Thus, the endogenous adenosine that tonically activates the A_1_R comes only from D_1_R positive MSNs in the medial thalamus-dorsomedial striatum circuit.

## Author contributions

SA: Conceptualization, Formal analysis, Funding acquisition, Investigation, Visualization, Methodology, Writing—original draft.

WTB: Conceptualization, Formal analysis, Resources, Supervision, Funding acquisition, Investigation, Visualization, Methodology, Writing—review and editing.

## Acknowledgements

This work was supported by R01DA042779 (WTB), ARCS Foundation (SA), and F30 DA051117 (SA). We thank Dr. John T. Williams for comments on this manuscript and financial support for this project R01DA008160 (JTW). We thank Dr. Brigitte Kieffer for providing us with Oprm1 fl/fl mice, Dr. Christopher Ford for providing Drd1-cre mice, and Dr. Tianyi Mao for providing Adora2a-cre mice. We also thank Ms. Katherine Suchland, Dr. James Bunzow, and Dr. Joe Lebowitz for genotyping the transgenic mice.

## Notes

**Conflict of Interest**: The authors declare that no competing interests exist.

### Competing Interest Statement

The authors have declared no competing interest.

